# Towards an understanding of injury in aortic tissue prior to actual failure: quantification of mechanically induced collagen damage using collagen hybridizing peptides

**DOI:** 10.64898/2026.11.18.767693

**Authors:** Anna Pukaluk, Daniel Ch. Haspinger, Heimo Wolinski, Christian Viertler, Janna C. Nawroth, Gerhard A. Holzapfel

**Author notes:** These authors jointly supervised this work (Janna C. Nawroth, Gerhard A. Holzapfel).

## Abstract

The development, validation, and optimization of endovascular aortic devices require quantitative assessment of tissue damage occurring during surgical interventions. Until now, this assessment—before complete failure—has been largely unfeasible for lack of a suitable methodology. Recently developed collagen hybridizing peptides (CHPs) allow selective detection of damage to collagen, the primary load-bearing constituent of the aortic wall. However, their application in large, mechanically loaded arteries has not yet been established. Therefore, we optimized an experimental workflow for large arteries that accounts for their energy-storage function and geometrical constraints; furthermore, we have characterized the tissue’s autofluorescence and selected a suitable CHP fluorophore. Specimens from a single porcine aorta were examined using uniaxial tensile testing, histological analysis, fluorescence spectral analysis, confocal microscopy, and quantitative image analysis to present collagen damage as a function of tissue stretch. The workflow detected load-induced collagen damage before tissue failure, whereas histological evaluation showed no obvious load-induced disruption. Damage scores were comparable between the undeformed control and the specimen stretched by a factor of 1.2, while specimens stretched beyond 1.2 exhibited elevated scores. This pattern corresponded to higher complication rates with excessive stent oversizing. Furthermore, the proposed protocol can be adapted for *ex vivo* stent deployment experiments to compare collagen damage resulting from various aortic device designs and deployment conditions. Quantitative damage metrics could inform structurally motivated constitutive material models for computational simulations of endovascular interventions. Combined with histological and structural analyses, this workflow could also help link tissue composition—or organization—to individual susceptibility to damage.

## 1 Introduction

Cardiovascular diseases are the leading cause of death worldwide [1]. Among these, aortic diseases—despite significant advances in diagnosis and treatment—remain associated with high mortality [2]. The modern treatment of numerous aortic diseases relies primarily on endovascular interventions. Aneurysms and dissections of the abdominal or thoracic aorta are typically managed via the endovascular implantation of stent grafts [3–5], whereas aortic isthmus stenosis (coarctation of the aorta) is treated using balloon angioplasty—either with [6] or without [7, 8] the implantation of a stent. Nevertheless, these life-saving procedures carry the risk of rare but potentially fatal complications [9], such as retrograde dissection [3, 4], the development of an aneurysm [7, 8], endoleaks [5], or recurrent aortic stenosis [6].

Endovascular treatments rely on the interaction between the aortic wall and a balloon or stent, which can lead to tissue damage as an adverse consequence [10, 11]. In particular, excessive oversizing of balloons or stents is considered a significant factor contributing to unfavorable clinical outcomes [3, 12, 13] (interestingly, such oversizing is even used intentionally in animal studies to induce traumatic stenosis in healthy arteries [14]). Both balloon angioplasty and the implantation of stents or stent grafts exert mechanical loading that causes the aortic wall to stretch; this can result in structural damage to individual tissue constituents or even rupture of the entire tissue.

From a mechanical perspective, collagen is of particular interest, as it constitutes the primary load-bearing component of the aortic wall [15]. It has been demonstrated that collagen in the aorta responds to mechanical loading through structural changes—at both the microscale [16, 17] and the nanoscale [18]. However, it remains unclear whether these structural changes are accompanied by damage to the collagen that precedes tissue failure. In this context, a recent study by Yusefi et al. [19] investigated the mechanical behavior of thoracic aortic tissue following endovascular stent grafting. Two distinct mechanical responses were observed: a stiffening of the distal aortic wall—suggesting the recruitment of collagen fibers—and a softening of the proximal aortic wall, which points to damage to the collagen fibers. The finding that mechanical loading of the tissue in the proximal aorta leads to collagen damage—going beyond mere fiber recruitment [19]—is consistent with clinical observations regarding aortic dissections. Such dissections frequently occur in the proximal aorta following thoracic endovascular aortic repair [3, 4]. These observations suggest that collagen damage—associated with device oversizing—could be a significant factor in life-threatening complications following endovascular repair.

Attempts have been made to identify collagen damage resulting from mechanical loading; however, this has proven difficult due to significant limitations in the available methodological approaches [20–23]. The examinations employed the gold standard for tissue assessment: histologically stained sections evaluated by experienced pathologists [20, 21]. While this approach allowed for the identification of tissue rupture, it lacked the sensitivity required to attribute damage to specific tissue components (such as collagen) or to detect damage at the level of individual collagen fibers. Furthermore, attempts were made to detect collagen damage using polarized light microscopy (PLM), with changes in the birefringence of the collagen serving as a damage marker [22]. Although PLM has been used in previous studies to assess thermal damage to collagen [24, 25], the results obtained were not validated against established markers for collagen damage—such as those derived from trypsin digestion; furthermore, some authors point out that the loss of birefringence is not specific to collagen but can occur in many tissue components, including, among others, elastin and smooth muscle [26]. Recently, Chakraborty et al. [23] used PLM to qualitatively investigate mechanically-induced collagen damage in the sclera; however, the results obtained proved unsuitable for quantitative analysis. It should be noted that signal intensity in PLM is influenced not only by collagen damage but also by fiber orientation [22]. Since the orientation of collagen fibers in aortic tissue changes under mechanical loading [27], such a reorientation of the fibers could be erroneously interpreted as collagen damage.

The limitations associated with the assessment of collagen damage—such as those encountered when analyzing histologically stained tissue sections or employing PLM—have been largely overcome through the development of collagen hybridizing peptides (CHPs) [28]. These are collagen-specific markers that have been validated via trypsin digestion and enable the quantification of collagen damage. CHP binds specifically to unfolded collagen molecules, while an attached fluorophore enables their visualization via fluorescence microscopy. Zitnay et al. [28] developed CHP to detect collagen damage at the molecular level, induced by macroscopic mechanical loading of rat tail tendons. The methodology was rapidly extended to further studies on mechanically-loaded tendons [29, 30] and subsequently applied to investigations of other collagenous tissues, such as bone [31], cranial meninges [32, 33], and cerebral arteries [34, 35]. Furthermore, it has been used to identify not only load-related but also disease-induced collagen damage [36, 37]. However, the use of CHP in arteries subject to mechanical loading has thus far been limited to small arteries—specifically, the sheep [34] and porcine cerebral arteries [35]. The aorta has only been studied in a mouse model and in the context of atherosclerosis development—without being subjected to mechanical loading [37]. To the best of our knowledge, the use of CHP to detect load-induced collagen damage in large arteries has not yet been investigated.

Applying the experimental protocol used for small arteries to large arteries is not straightforward. First, the use of CHP in large arteries—which are subject to mechanical loads—might be limited by the fact that collagen damage can arise through various mechanisms; a phenomenon already documented in tendons with different physiological functions [29]. While CHP proved effective for investigating collagen damage in positional tendons (those serving to stabilize posture), its application to energy-storing tendons met with only limited success [29]. Since energy storage—unlike in small arteries—is a central physiological function of large arteries [38], the applicability of CHP cannot be assumed without further evidence. Second, small arteries (or the large arteries of small mammals) are thin enough to be successfully stained with CHP and imaged in their intact state [34, 35, 37]; however, for the thick-walled aorta, the methods for reliably preparing sections and subsequently imaging them must be adapted accordingly. Here, we present an experimental workflow for applying CHP to a mechanically loaded porcine aorta—a model frequently used as an analogue for the human aorta [39–41].

We demonstrate the feasibility of using CHP to detect load-induced collagen damage in aortic tissue—well before mechanical failure occurs. The proposed protocol could support the experimental assessment of tissue damage caused by aortic devices [19]. The resulting quantitative damage measurements could be integrated into structurally motivated constitutive material models [42] that account for collagen damage when predicting the mechanical behavior of aortic tissue. Furthermore, this approach—combined with detailed histological and structural investigations [43]—could help explore how tissue structure influences susceptibility to damage.

## 2 Methods

### 2.1 Aortic specimen

The experimental workflow was applied to a descending aorta obtained from a domestic pig (*Sus domesticus*, approximately six months old, body weight approximately 100 kg) from a local slaughterhouse. Porcine aortic tissue was selected to ensure that all specimens consisted of healthy tissue; this is because adverse clinical events—such as retrograde dissection—typically occur in aortic tissue that was healthy prior to the intervention [3]. Furthermore, the use of aortic tissue from young pigs avoids age-related pathological changes, such as atherosclerosis. Since atherosclerosis itself can lead to collagen damage [37], it could confound the identification of load-induced damage. In contrast, human aortic tissue obtained during autopsies is frequently affected by atherosclerosis—at least to some extent [16, 44].

The aorta was transported to the laboratory on ice within 1 h of slaughter and processed further within 60 h. To prevent tissue dehydration, the tissue was moistened with phosphate-buffered saline (PBS; pH 7.4) during handling at room temperature and stored in PBS at 4*^◦^*C between the individual processing steps. After removing the surrounding loose connective tissue, the aortic arch was excised, and the descending aorta was cut open longitudinally (Fig. 1(a)). The intimal and adventitial layers were carefully dissected away from the media layer [45]. The medial layer was selected because it acts as the primary load-bearing layer of the aortic wall under physiological loading conditions [46]; furthermore, it is the primary site for tissue injury associated with the non-physiological mechanical loading during endovascular interventions [10, 11].

**Figure 1:**
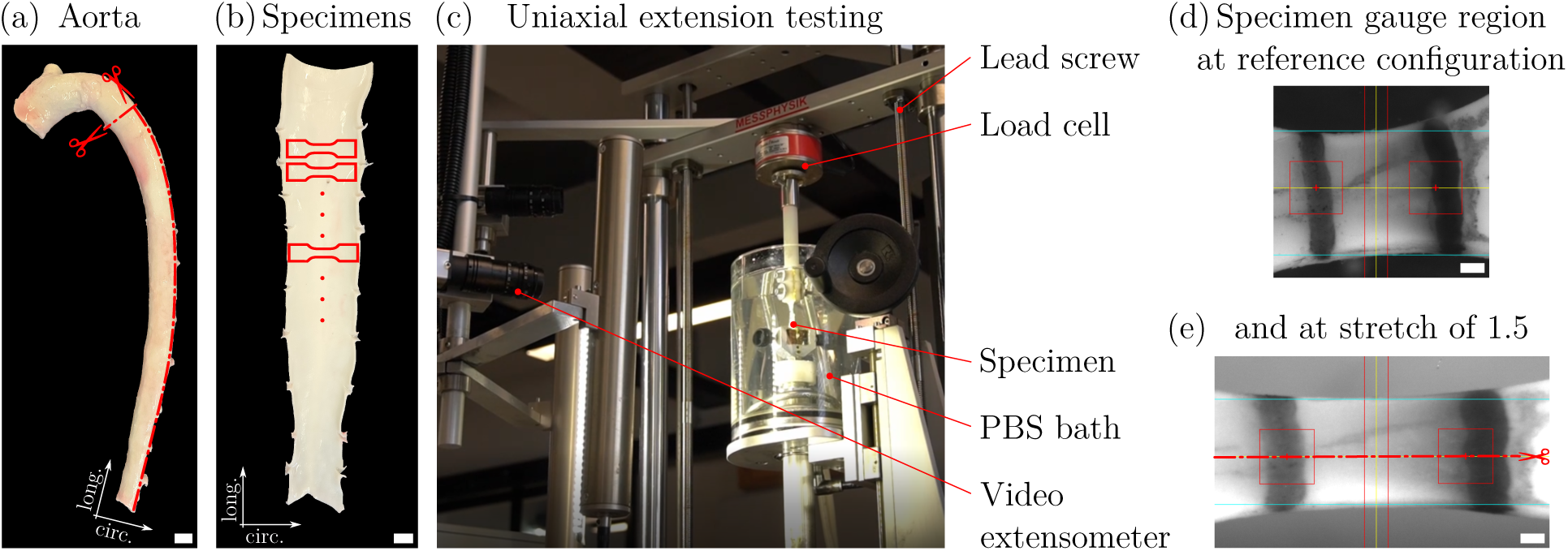
Sample preparation and mechanical testing, consisting of (a) the excision of the aortic arch and a longitudinal opening of the descending aorta (indicated by the red dashed paths marked with scissors symbols), (b) the cutting out of dog-bone-shaped specimens (solid red areas) [47] following the preparation of the medial layer (not shown), and (c) the mounting of the specimen into the testing device [48] for uniaxial extension testing. Representative tracking of the gauge region is shown for a specimen in its reference configuration (d) and in a stretched state at a stretch of 1.5 (e). After fixation, the stretched specimen was cut into two halves (red dashed line, marked with a scissors symbol) to allow for histological staining both within the plane and perpendicular to it. The anatomical directions of the aorta are indicated by arrows (longitudinal, long.; and circumferential, circ.). The scale bars correspond to 1 cm in (a) and (b), and 1 mm in (d) and (e).

### 2.2 Uniaxial extension testing

For the mechanical tests, dog-bone-shaped specimens were excised from the aortic media and oriented in the circumferential direction (*n* = 7) (Fig. 1(b)). The dog-bone-shaped geometry was chosen to ensure uniform deformation within the gauge region [49]; this region was marked by two dots applied using a tissue marker, spaced approximately 5 mm apart [47]. The circumferential direction was selected because it corresponds to the primary loading direction to which the aortic wall is subjected during balloon angioplasty or stent deployment. The reference thickness *T* of each specimen was determined optically [45].

The uniaxial extension tests were conducted using a multi-purpose testing device [48] (Fig. 1(c)). In brief, the specimen extension was controlled via a stepper motor that drove lead screws, which converted the rotational motion into linear displacement. The axial force *f* acting on the specimen was measured using a load cell. The reference width *W* and length *L* of the undeformed specimen were measured at the time of clamping; the current specimen width *w* and length *l* were recorded during the test using a video extensometer—details regarding the technical specifications can be found in [48]. To replicate the *in vivo* environment, the specimens were kept in PBS at 37*^◦^*C during the test.

The specimens were tested using a stretch-controlled protocol based on measurements from a video extensometer (Fig. 1(d) and (e)). The stretch rate was set to 5 mm*/*min, following established protocols [21, 50], as the stretch rate occurring during balloon angioplasty or stent deployment is not precisely defined. Each specimen was subjected to five preconditioning cycles to achieve reproducible mechanical behavior; this approach is based on a preliminary investigation as well as previous studies, which indicate that five [47, 51] or even three [21, 50] preconditioning cycles are sufficient. The preconditioning was performed up to a stretch of 1.2, which roughly corresponds to the physiological stretch experienced during the cardiac cycle [52–54]. After the preconditioning, the specimens were subjected to a loading and unloading cycle up to a predefined maximum stretch. The target stretch differed between specimens, ranging from 1.2 to 1.8 in increments of 0.1. After the experiment, each specimen was stretched again to its respective, previously determined maximum stretch. While held in this stretched state, the specimen was immersed in a 4% formaldehyde solution for 4 h at room temperature to fix the state of deformation for subsequent analysis of collagen damage.

The mechanical behavior under uniaxial extension was quantified using the Cauchy stress, which was calculated under the assumption of incompressibility as

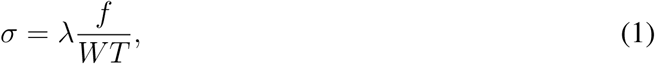

where *λ* = *l/L* is the stretch, derived from the reference gauge length *L* and the current gauge length *l*, both measured using a video extensometer; *W* is the reference width, *T* is the reference thickness, and *f* is the force measured during the test by means of a load cell.

### 2.3 Histological staining

Each mechanically tested specimen was fixed in formaldehyde and bisected in the circumferential direction (the direction of loading) (Fig. 1(e)). In addition, a rectangular, undeformed reference specimen was taken from the aortic media to serve as a control for comparison with the mechanically loaded specimens. This reference specimen was also bisected. During paraffin embedding, one half of each specimen was oriented in-plane, showing the circumferential-longitudinal plane, while the other half was oriented out-of-plane, showing the circumferential-radial plane (Fig. 2). Each paraffin-embedded piece of the aortic media was sectioned into slices approximately 4 µm thick using a Microm HM 335 microtome (Microm, Walldorf/Baden, Germany).

**Figure 2:**
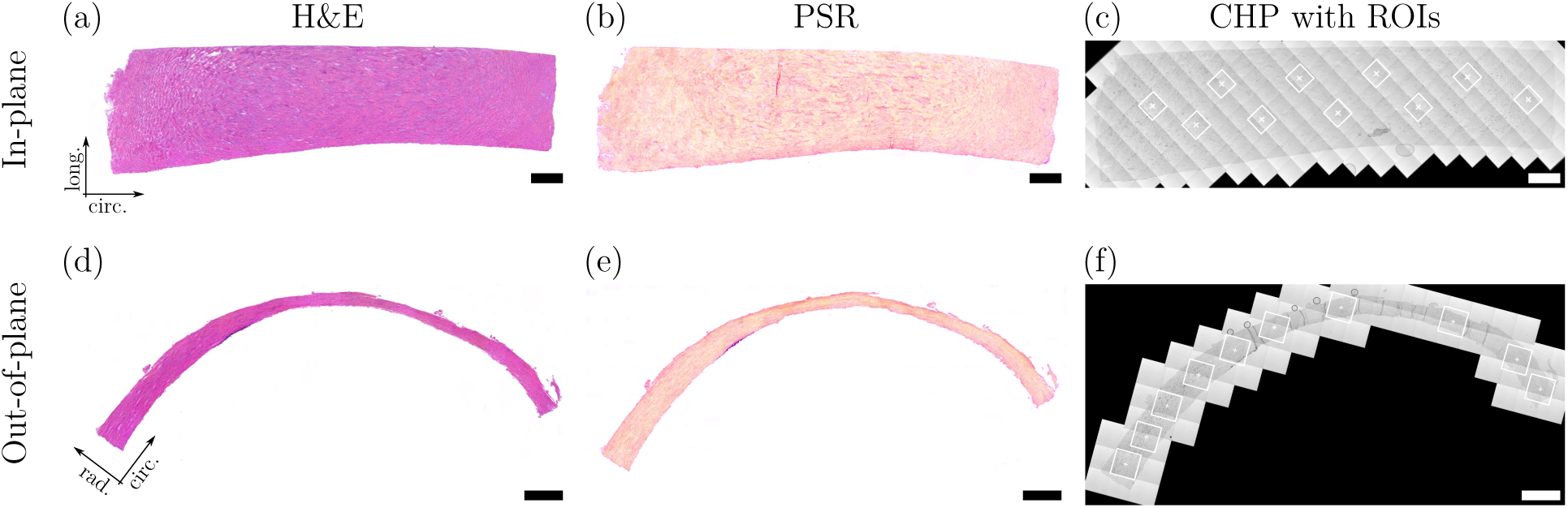
Representative histological images of a specimen stretched to 1.5 times its original length; shown are orientations within the plane (a)–(c) and out-of-plane (d)–(f). The sections were stained with hematoxylin and eosin (H&E), (a),(d), and with picrosirius red (PSR), (b),(e). Transmitted-light mosaic scans of the sections stained with collagen hybridizing peptides (CHP) show, in (c) and (f), the locations of the regions of interest (ROIs) examined, which are marked by white squares. The anatomical directions of the aorta are indicated by arrows (longitudinal, long.; circumferential, circ.; and radial, rad.). The scale bars correspond to 1 mm.

Adjacent sections of each paraffin-embedded sample were stained with hematoxylin and eosin (H&E) for routine tissue assessment, with picrosirius red (PSR) to visualize fibrillar collagen, and with CHP to detect mechanically induced collagen damage (Fig. 2). Additional sections of the reference specimen were deparaffinized—but not stained—and used for spectral scanning to identify a fluorophore that exhibits minimal spectral overlap with the tissue’s autofluorescence (Section 2.4).

CHP staining was performed according to the 3Helix protocol [36, 55] using F-CHP, which had been selected beforehand based on spectral scans (Sections 2.4 and 3.3). In brief, CHP powder was dissolved in PBS to prepare a 100 µM stock solution, which was subsequently diluted with PBS to a concentration of 20 µM. Immediately prior to staining, the diluted CHP solution was heated to 80*^◦^*C for 5 min to dissociate the CHP triple helices into single strands. The solution was then rapidly cooled to room temperature before being applied to the tissue sections. The CHP-stained sections were incubated overnight at 4*^◦^*C and subsequently washed three times in PBS for 5 min each at room temperature.

### 2.4 Confocal microscopy

Imaging was performed at IMB-Graz Optical Imaging Resource (University of Graz, Austria) using a Leica SP8 confocal microscope (Leica Microsystems, Mannheim, Germany) equipped with an argon laser and a Fluotar VISIR 25*×* 0.95 NA water-immersion objective.

First, spectral scans were performed to select a suitable CHP conjugate whose emission spectrum does not overlap with the broad autofluorescence spectrum of the aortic media [16]. Although the 5-FAM conjugate (F-CHP) is the most commonly used [29, 30, 33–35], the Cy3 conjugate (R-CHP) is recommended for samples with strong autofluorescence [36, 55]. Other conjugates, including biotin (B-CHP) [36], are also available. Spectral scans were performed on diluted solutions of F-CHP and R-CHP, on intact aortic media, and on formaldehyde-fixed, paraffin-embedded sections (following deparaffinization). The intact aortic media—that is, the medial layer obtained solely through dissection, without any further treatment—was investigated as a potential alternative to histologically processed medial sections. F-CHP and R-CHP solutions were excited at 488 nm and 514 nm, respectively; emission spectra were recorded in the ranges of 495–645 nm and 520–695 nm, respectively. Intact preparations of the aortic media, as well as deparaffinized sections of the media, were examined using the excitation and emission parameters specific to F-CHP and R-CHP. In each experimental setup, the emission was recorded using photomultiplier tube detectors with a constant bandwidth of 5 nm. The captured images had a size of 512 *×* 512 pixels, with a pixel size of 1.2 µm and a bit depth of 8.

Following the selection of F-CHP as the CHP conjugate (Results, Section 3.3), images were acquired using excitation at 488 nm and hybrid detectors in photon-counting mode. One detection channel recorded F-CHP fluorescence (525–540 nm)—an indicator of collagen damage—while a second channel recorded tissue autofluorescence (565–580 nm) to allow for a more accurate estimation of the total tissue area. Ten regions of interest (ROIs) were imaged on each section. For each of these regions, a *z*-stack was acquired—with an image size of 1024 *×* 1024 pixels, a pixel size of 0.6 µm, and a bit depth of 12. The step size for the *z*-stack was 3 µm. In addition, a mosaic scan was performed using transmitted light to record an overview image of each section, which also includes the positions of the ROIs (Fig. 2(c),(f)).

### 2.5 Image processing

Image processing was performed according to Anderl et al. [35], with modifications made to adapt it to the specific sectioning protocol and the imaging setup (Fig. 3). From each *z*-stack, the image slice with the highest integrated F-CHP fluorescence intensity was selected as representative of the collagen damage within the specimen [35]; this accounts for the fact that the cut surfaces were uneven. Alternative approaches include selecting the middle slice of the *z*-stack [33] or using a *z*-projection [31]. However, collagen damage is not necessarily located in the middle slice, and a *z*-projection can complicate the assessment of damage in sections with non-uniform thickness. Background pixels were excluded by masking the F-CHP image using the corresponding autofluorescence image. Only pixels with a non-zero autofluorescence intensity were retained, ensuring that only tissue-covered areas were included in the analysis. The use of hybrid detectors in photon-counting mode—characterized by low detector noise—enables reliable, intensity-based masking. Consequently, an automated procedure was preferred over the manual masking method described by Anderl et al. [35], as it was deemed more efficient and reproducible for the workflow in question.

**Figure 3:**
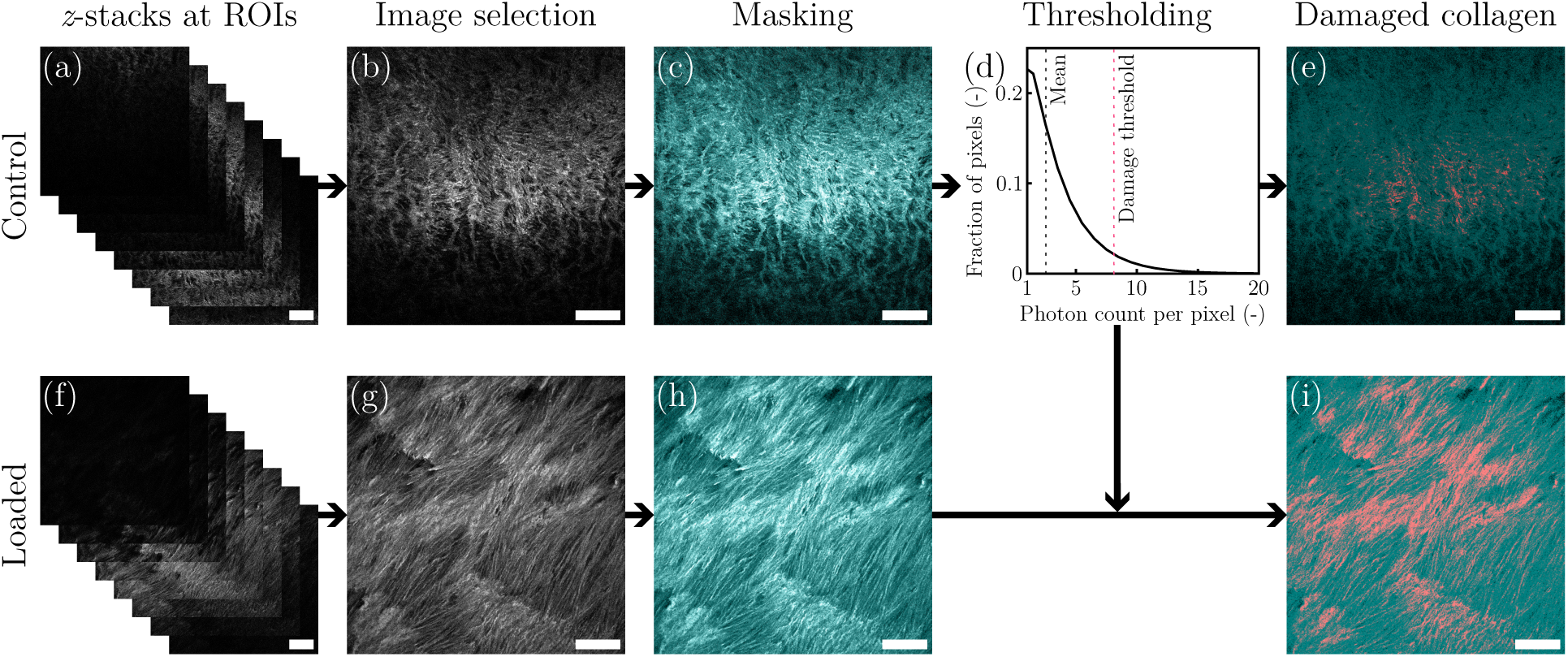
Image processing workflow, illustrated using representative *z*-stacks and images of a control specimen (undeformed) (a)–(e) and a loaded specimen that has been stretched to 1.5 times its original length (f)–(i). From each *z*-stack (a),(f), the image with the highest F-CHP fluorescence intensity was selected (b),(g) and masked using the corresponding autofluorescence image—shown in cyan—(c),(h). The damage threshold was defined determined on all ROIs from the control specimen (d) and was used to identify damaged collagen (shown in red) in both the control (e) and the loaded specimen (i). The scale bars correspond to 100 µm.

The damage threshold was defined as the mean intensity plus two standard deviations, calculated across all ROIs of the undeformed reference, which served as a control [35]. This threshold restricted the detection of collagen damage to those fluorescence outliers that exhibited the highest intensity. For each mechanically loaded section, pixels with a fluorescence intensity below the damage threshold were excluded; the proportion of pixels that exceeded this threshold served as a quantitative measure of collagen damage [35].

## 3 Results

### 3.1 Mechanical response

Under uniaxial extension, the aortic media exhibited the characteristic mechanical behavior (Fig. 4). The loading and unloading curves did not coincide, which confirmed the expected hysteresis [56]. Nonlinear stiffening—typically attributed to the involvement of collagen fibers [56]—occurred at higher stretch levels and was particularly pronounced in the specimen stretched by a factor of 1.8. Since the aortic media typically fails at a stretch of approximately 2 [47], the experiments covered the mechanical behavior from the physiological loading range up to the point immediately preceding tissue failure. Minor differences between individual samples likely reflect regional variations in the mechanical properties of the aortic wall [57, 58]—especially given that the use of samples from a single aorta ruled out differences between different animals.

**Figure 4:**
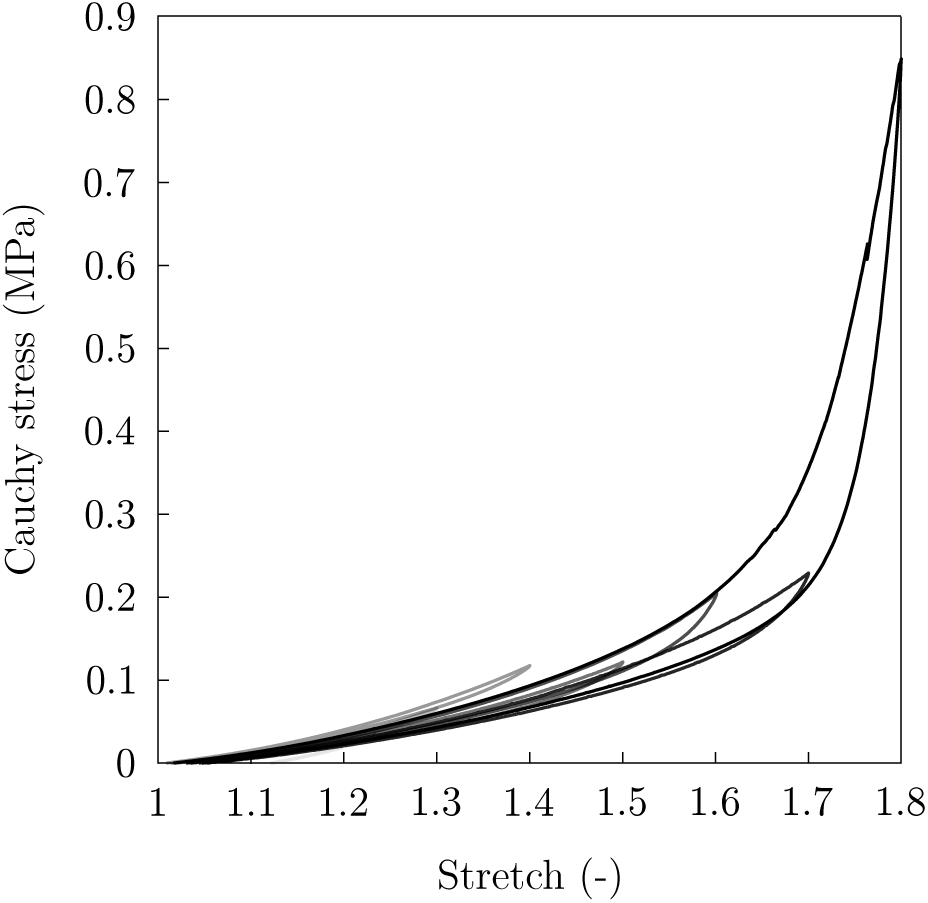
Cauchy stress-stretch behavior of the media of the porcine aorta under uniaxial extension in the circumferential direction up to various maximum stretch levels.

### 3.2 Histological investigation

The H&E- and PSR-stained sections showed no apparent disruption of tissue integrity resulting from mechanical loading; this is consistent with the fact that no tissue failure occurred in the specimens during mechanical testing. Visible tissue distortions, such as lamellar separation (Fig. 2), were attributed to artifacts resulting from the sectioning process on the microtome—caused by differences in the mechanical properties of adjacent lamellae; these were also observed in the undeformed control sections.

### 3.3 Fluorescence spectra

F-CHP and R-CHP showed relatively narrow emission spectra, with emission maxima at approximately 530–535 nm (excitation at 488 nm) and 560–565 nm (excitation at 514 nm), respectively (Fig. 5). In contrast, the intact aortic media exhibited broad autofluorescence spectra, with an emission maximum at approximately 535 nm under 488 nm excitation (Fig. 5(a)) and at approximately 560 nm under 514 nm excitation (Fig. 5(b)). The autofluorescence remained at a high level within the ranges of 495–610 nm and 520–620 nm, respectively. The overlap between the CHP emission spectra and the autofluorescence spectrum of the intact aortic media suggested that none of the conjugates would be suitable—unless the CHP fluorescence intensity was sufficiently higher than the tissue’s autofluorescence to allow for reliable signal discrimination; however, this was not the case in our preliminary studies (data not shown). In contrast, sections of the aortic media—when excited with both 488 nm and 514 nm—exhibited relatively narrow spectra with an emission maximum at approximately 570 nm. This emission maximum overlapped with the R-CHP emission spectrum but differed significantly from that of F-CHP, indicating minimal spectral overlap between F-CHP fluorescence and tissue autofluorescence. Therefore, F-CHP was selected as the CHP conjugate for all subsequent experiments.

**Figure 5:**
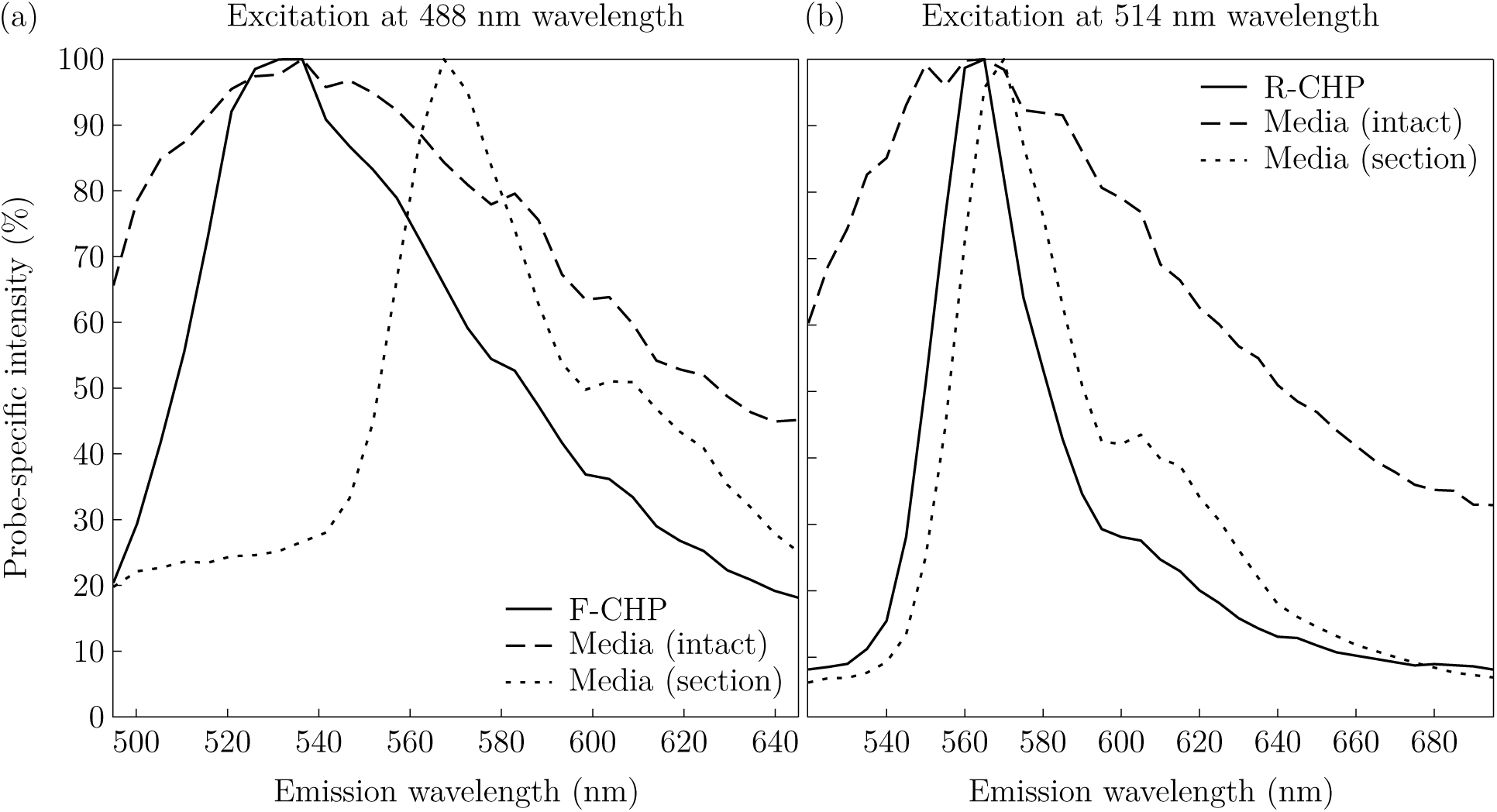
Emission spectra recorded with (a) 488 nm excitation and (b) 514 nm excitation. Each panel shows the corresponding CHP conjugate (F-CHP or R-CHP) as well as the spectra of the aortic media—both after dissection alone (intact) and following histological processing (section).

### 3.4 Collagen damage evaluation

For each specimen, both in-plane and out-of-plane sections were evaluated. However, the out-of-plane sections proved unsuitable for a quantitative assessment of collagen damage and were therefore excluded from further analysis (data not shown). This observation is consistent with a recent study showing that images of aortic tissue acquired out-of-plane are fundamentally unsuitable for the quantification of collagen—particularly when high axial resolution is employed [59], such as that achieved through techniques like second-harmonic generation or confocal microscopy. Consequently, further analyses were performed using in-plane sections, adopting an approach from previous studies on cerebral arteries [34, 35].

The percentage of damaged collagen was similar for the undeformed control specimen (stretch equal 1.0) and the specimen stretched to a value of 1.2; in both cases, the damage scores for the individual ROIs were approximately 4% (Fig. 6). For specimens stretched beyond a value of 1.2, the percentage of damaged collagen tended to increase with greater stretch, while the damage scores varied significantly more between the individual ROIs within each respective specimen.

**Figure 6:**
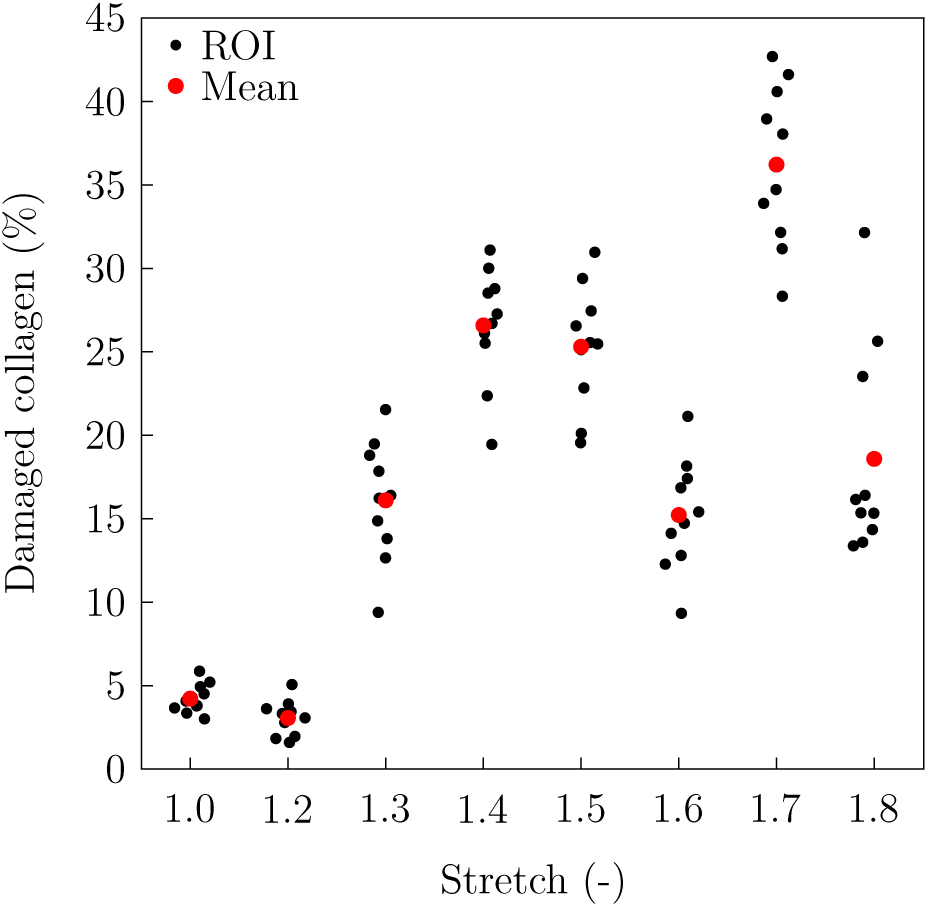
Percentage of collagen damage, measured in ten ROIs for each stretch level (black), with the corresponding mean values (red).

## 4 Discussion

### 4.1 Feasibility of using CHP for damage detection in aortic tissue prior to failure

Identifying load-induced collagen damage in aortic tissue represents a step toward understanding how aortic devices and biomaterials contribute to clinical complications following endovascular interventions. In the present study, CHP was used to detect collagen damage that precedes structural failure; in contrast, routine histological examinations showed no obvious, load-related disruption (Fig. 7). Such damage could contribute to complications following endovascular interventions (Section 4.2). Quantifying this damage could therefore support experimental and numerical studies on how mechanical loading affects the behavior of collagen and aortic tissue (Section 4.3), as well as investigations into the tissue’s susceptibility to damage (Section 4.4).

**Figure 7:**
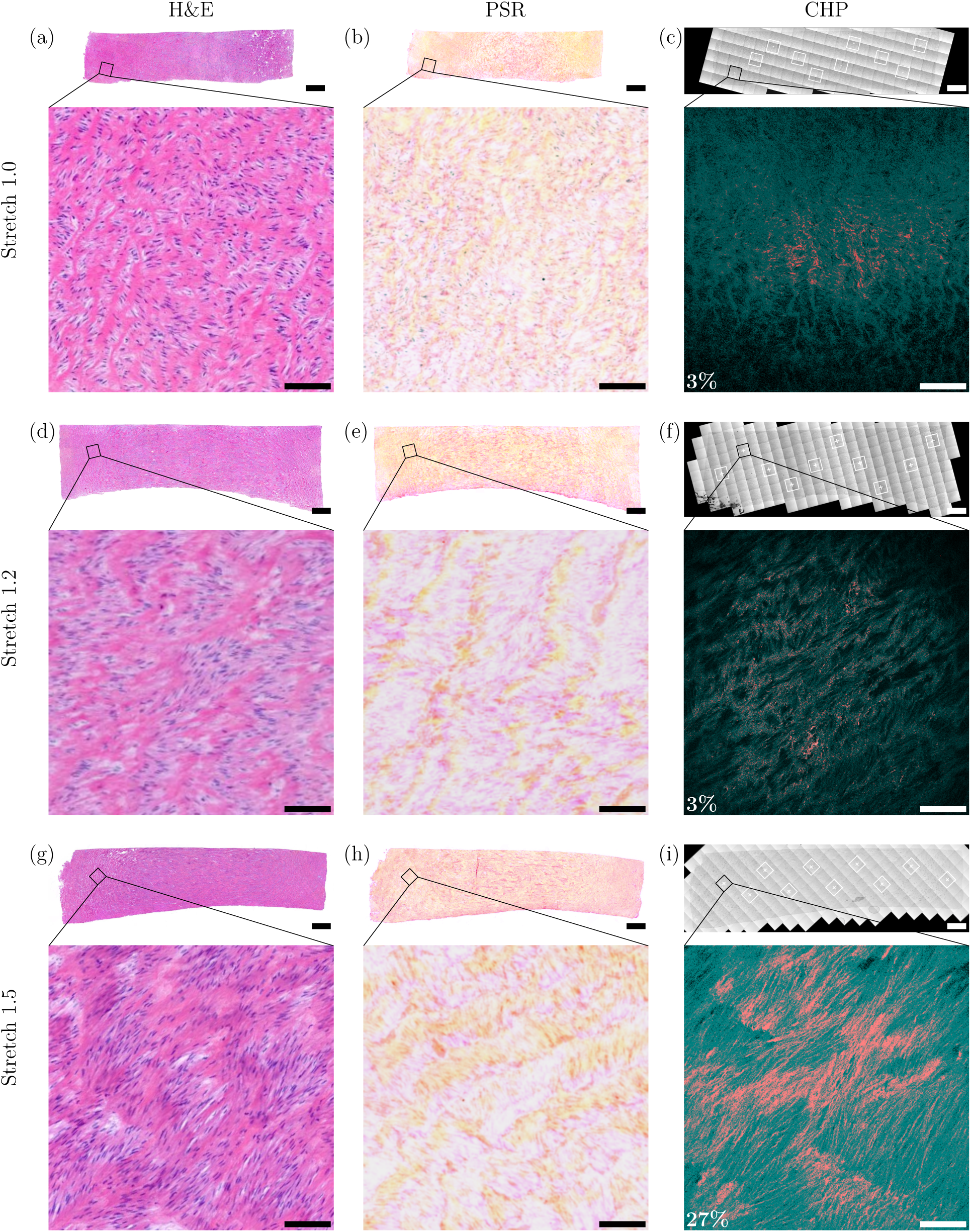
Representative histological images and CHP images of the control specimen (stretch 1.0) – (a)– (c) – as well as of the specimens stretched to 1.2 – (d)–(f) – and 1.5 – (g)–(i), respectively. The sections were stained with hematoxylin and eosin (H&E) (a),(d),(g), picrosirius red (PSR) (b),(e),(h), and collagen hybridizing peptides (CHP) (c),(f),(i). Black squares in the overview images indicate the areas shown in the corresponding detailed images. White squares in the CHP overview images—which are transmitted-light mosaic scans (c),(f),(i)—mark the locations of the additional ROIs that were examined. In the CHP close-up images, the autofluorescence of the tissue is shown in cyan, and the damaged collagen is shown in red. For each of the ROIs displayed, the percentage of collagen damage is indicated. The scale bars correspond to 1 mm in the overview images and 100 µm in the close-up images.

Our results show that load-induced collagen damage in large arteries can be detected using CHP—regardless of their energy-storage function [29]. These findings suggest that load-induced collagen damage is accompanied by the unfolding of collagen triple helices, which is consistent with the detection mechanism proposed for CHP [28]. However, the experimental workflow had to be adapted to the relatively thick aortic tissue. Interestingly, the sample preparation—consisting of formaldehyde fixation, paraffin embedding, deparaffinization, and subsequent tissue sectioning—significantly altered the autofluorescence spectrum of the aortic media. The emission spectra presented in this work facilitated the targeted selection of fluorophores and can serve as a guide for future studies when choosing appropriate fluorophores for either intact specimens or processed specimens that have been sectioned.

The quantified collagen damage provides insights into the relationship between collagen damage and the applied stretch. The fact that no increased collagen damage occurs at a stretch of 1.2 compared to the undeformed control suggests that this level of deformation can be reached without causing additional collagen damage. The increase in collagen damage at a stretch greater than 1.2 suggests a relationship between the extent of tissue deformation and collagen damage. Finally, the greater variability in damage values within individual specimens stretched beyond a value of 1.2 points to a heterogeneous distribution of the damage.

Unexpectedly, the control specimen already exhibited a degree of collagen damage of approximately 4%; a result that was also observed in the specimen subjected to a stretch of 1.2. The normal turnover of collagen fibers has a half-life of approximately 70 days, with an estimated degradation rate of 1–2% per day [60] – a figure that is lower than the extent of collagen damage we observed in our control sample. Since the control specimen was not subjected to any mechanical loading, the observed damage is presumably attributable to another cause. Non-specific binding of CHP is unlikely, given its high specificity for unfolded collagen and the absence of binding sites within helically-folded collagen molecules [61]. The neutral and hydrophilic nature of CHP further reduces the likelihood of non-specific binding [61]. However, a certain degree of collagen denaturation may have been caused by the sample preparation process—including formaldehyde fixation, paraffin embedding, and deparaffinization [61, 62]. The inclusion of an undeformed control sample is therefore crucial for establishing a baseline for collagen damage that is not attributable to mechanical loading.

### 4.2 Relevance of stent oversizing for clinical practice

Although the present study was conducted using tissue from a single aorta, the observed relationship between collagen damage and applied stretch is consistent with clinical observations regarding stent oversizing. In particular, the absence of detectable collagen damage up to a stretch of 1.2 aligns with clinical findings that moderate stent oversizing is generally well tolerated, whereas more extensive oversizing is associated with higher complication rates [3, 12, 63]. Consistent with our observation that a comparable degree of collagen damage occurs at stretch ratios of 1.0 and 1.2, Xiang et al. [63] reported no differences in adverse clinical outcomes among patients undergoing thoracic endovascular aortic repair—with stent oversizing of approximately 2% (range 0%–5%) and 10% (range 5%–25%), respectively. However, in patients who underwent thoracic endovascular aortic repair (TEVAR) with a stent oversizing of more than 20%—as observed by Kpodonu et al. [3] and Canaud et al. [12]—retrograde aortic dissection occurred more frequently. Liu et al. [13] proposed a somewhat lower threshold; they reported fewer cases of retrograde aortic dissection in patients treated with an average stent oversizing of approximately 1% (range 0%–5%), compared to patients in whom an oversizing of approximately 19% (range 8%–32%) was applied.

The broad time frame within which adverse events can occur following endovascular interventions may be attributable to varying biological responses to collagen damage. Adverse events such as aortic dissection—which may occur either immediately [3, 4] or within a few days of the procedure [4, 13]—could reflect acute tissue damage caused by mechanical overload. However, adverse events—such as a dissection [3, 4, 12], an aneurysm [7, 8], or a stenosis [7]—can also occur months or even years later. These delayed complications may still be attributable to the mechanical loading exerted during the procedure. It has been established that mechanical stress can induce structural changes in the collagen and elastin within the aorta [16, 17]. Such structural alterations are characteristic of extracellular matrix remodeling—a process known to drive the progression of disease [64, 65].

### 4.3 Potential for the preclinical evaluation of aortic devices

The proposed experimental workflow offers significant potential for assessing collagen damage caused by aortic devices. Although the method was originally developed using uniaxial extension tests, it can be adapted to other mechanical loading conditions and integrated into *ex vivo* experiments—such as the experimental setup proposed by Yusefi et al. [19], in which an aorta is subjected to a physiologically relevant load followed by stent deployment. Collagen damage could then be compared across various experiments that involved differing degrees of stent oversizing, varying device geometries or materials, inflation pressures, or other procedure-related variables.

Furthermore, the quantitative scores for collagen damage determined via the proposed workflow could support *in silico* simulations of stent deployment. These experimental data could serve as a basis for structurally motivated constitutive material models [42] by assisting in the calibration of damage-related parameters or in the evaluation of model predictions. Subsequently, these models could be employed in simulations of aortic stent-graft deployment, where predictions regarding device-induced wall stresses [66] would be complemented with predictions concerning collagen damage. Taken together, these *ex vivo* and *in silico* applications could facilitate the preclinical optimization of endovascular devices and their associated deployment strategies.

### 4.4 Further prospective applications

The experimental workflow was developed using porcine aorta, as it exhibits physiological similarities to the human aorta [58] and is frequently employed in the realistic testing of surgical procedures and devices intended for clinical use [39–41].

Nevertheless, extending the workflow to human tissue remains an important future goal. However, before the workflow can be applied to human tissue, various challenges must be overcome. Many age-related diseases are accompanied by collagen degeneration [36], which must be distinguished from collagen damage caused by mechanical stress. In particular, it has been shown that atherosclerosis leads to collagen damage [37]. Notably, atherosclerotic lesions—which are frequently encountered in autopsy samples [16, 44]—are not uniformly distributed [44, 67, 68]. This requires the careful selection of an appropriate control specimen to establish a baseline value for collagen damage not attributable to mechanical stress. Furthermore, aortic diseases are frequently accompanied by calcification that exhibits strong autofluorescence [69, 70]. This necessitates a careful selection of the conjugated fluorophore to avoid overlaps in the emission spectra.

On the other hand, the variability of human tissue offers the opportunity to investigate differences in susceptibility to damage. It is well established that aortic diseases affect both the mechanical behavior of aortic tissue [68, 71, 72] and the structural organization of its components [73–75]. The combination of histological and structural examinations [43] and quantitative measurements of load-induced collagen damage—under comparable loading conditions—could help clarify whether differences in tissue composition and collagen organization are associated with specific damage responses. Such studies could ultimately support the development of models that allow for the prediction of an individual’s susceptibility to injury or failure of the aortic wall.

The experimental workflow is not limited to aortic tissue; it can readily be applied to other large arteries, such as peripheral arteries. Endovascular interventions—including balloon angioplasties and stent placement—are also frequently performed in these vessels. Furthermore, the oversizing of stents is considered a critical factor that can contribute to unfavorable outcomes following such interventions [76, 77]. Consequently, the proposed workflow could provide a general framework for investigating mechanically induced collagen damage in various large arteries across different species.

## 5 Conclusion

Potentially fatal complications following endovascular interventions are thought to be caused by excessive mechanical loading of the aortic wall, resulting from the use of an oversized stent. However, the underlying mechanisms remain largely poorly understood. This study investigated load-induced collagen damage that could contribute to these complications. We have demonstrated the feasibility of using CHP to detect load-induced collagen damage in the large arteries of large mammals and have established an experimental workflow for its application. The observed relationship between collagen damage and applied stretch is consistent with clinical observations regarding the oversizing of stents. Mechanical loading within the physiological range did not result in any detectable additional collagen damage, whereas collagen damage increased at stretch levels that exceeded the physiological range. The proposed experimental workflow can be applied to both *ex vivo* and *in silico* studies, providing quantitative data to support the development, validation, and optimization of endovascular interventions and devices. Finally, the workflow is not limited to the porcine aorta; it can easily be extended to other large arteries in various animal models and, ultimately, to human tissue.

## Acknowledgements

The authors thank Sylvia Eidenhammer from the Medical University of Graz, Austria, for performing the CHP staining, as well as Jeff Weiss, Jared Zitnay, Allen Lin, and Lucas H. Timmins from the University of Utah, USA, for practical advice on working with CHP. The authors also thank Inge Fourneau from KU Leuven, Belgium, for an insightful discussion regarding the clinical aspects of stent oversizing. A. Pukaluk acknowledges the Alexander von Humboldt Foundation for its support (Humboldt Research Fellowship). G.A. Holzapfel acknowledges the Austrian Science Fund (FWF; Grant DOI: 10.55776/ I4545) for its support.

